# Phytohormone inhibitor treatments phenocopy brassinosteroid and gibberellin dwarf mutant interactions in maize

**DOI:** 10.1101/121772

**Authors:** Norman B. Best, Guri Johal, Brian P. Dilkes

## Abstract

Phytohormone biosynthesis produces metabolites with profound effects on plant growth and development. Modulation of hormone levels during developmental events, in response to the environment, by genetic polymorphism, or by chemical application can reveal the plant processes most responsive to a phytohormone. In many cases, chemical inhibitors are applied and the levels of specific phytohormones are measured to determine if, and which, phytohormone is affected by a molecule. In many cases, the sensitivity of biochemical testing has determined multiple pathways affected by a single inhibitor. Genetic studies are not subject to this problem, and a wealth of data about the morphological impacts of hormone biosynthetic inhibition has accumulated through the study of enzyme mutants. We previously identified a complex interplay between brassinosteroid (BR) and gibberellin (GA) in maize, where the interdependence of the two differs dependent on the developmental context. We found that: GA is required for loss of BR to induce retained pistils in the tassel florets (POPIT); BR is required for the loss of GA to induce tiller outgrowth; BR and GA are additive for plant height; BR has no effect on the induction of anther retention in ear florets of GA mutants. In this work, we sought to assess the specificity of three triazole inhibitors of cytochrome P450s by determining their abilities to recapitulate the phenotype of double mutants. The GA biosynthetic inhibitors uniconazole (UCZ) and paclobutrazol (PAC) were applied to the BR biosynthetic mutant *na2* and all double mutant phenotypes were recovered in the UCZ treatment. PAC was unable to suppress the retention of pistils in the tassels of *nana plant2* (*na2*) mutant plants. The BR biosynthetic inhibitor propiconazole (PCZ) suppressed tiller outgrowth in the GA biosynthetic mutant *dwarf5* (*d5*). All treatments were additive with genetic mutants for effects on plant height. Due to additional measurements done here but not in previous studies of the double mutants, we detected new interactions between GA and BR biosynthesis affecting plastochron index and tassel branching. These experiments, a refinement of our previous model, and a discussion of the extension of this type of work are presented.

## Introduction

Phytohormones control plant growth and development at very low concentrations. The entire life cycle of plants are influenced by the availability and amount of these metabolites. The number of identified classes of phytohormones currently includes auxins, cytokinins (CK), abscisic acid (ABA), jasmonic acid, salicylic acid, strigolactones, brassinosteroids (BR), and gibberellins (GA).^1–3^ The biosynthesis, transport, signaling, and responses to these phytohormones have been intensely studied in many species, including *Zea mays* (maize). Both biochemistry and genetic studies identified genes encoding phytohormone biosynthetic steps and mutants blocking hormone biosynthesis in many genetic model systems including *Pisum sativum* (pea), *Solanum lycopersicum* (tomato), *Arabidopsis thaliana* (Arabidopsis), and maize.^4–8^ Despite these studies, the molecular identities of many steps are as yet unidentified. Furthermore, the roles of phytohormones in processes that do not occur in established genetic models are less well explored. Biochemical inhibitors, particularly the triazoles, have elucidated physiological effects of these phytohormones in plant systems that lack the expansive genetic resources of species with vibrant genetics research communities like Arabidopsis and maize. The utility of inhibitor treatments for determining the roles of phytohormones in plant processes is determined by the specificity of the biochemical inhibitors. A test of this specificity is available if we determine the concordance between phenotypes induced by reducing hormone biosynthesis via genetic ablation of biosynthetic enzymes and the phenotypes observed after chemical inhibition.

A number of classical maize dwarf mutants are now demonstrated to encode GA and BR biosynthetic genes. Many mutants of maize affecting steps in gibberellin biosynthesis have been identified, including *ent*-copalyl synthase (*anther ear1*), *ent*-kaurene synthase (*dwarf5; d5*), CYP88A3 (*d3*), and GA3oxidase (*d1*).^9–13^ In addition to reduced stature, GA biosynthetic mutants of maize exhibit retained anthers in the normally pistillate ear florets, increased tiller outgrowth, shortened and broadened leaves, and decreased primary tassel branching.^10^ Maize mutants in three steps of BR biosynthesis are known, including disruptions of a Δ24-sterol reductase (*nana plant2; na2*), 5α-steroid reductase (*na1*), and CYP85A1/BR-6-oxidase (*brassinosteroid deficient1; brd1*).^8,14–17^ In addition to dwarfism, the maize BR biosynthetic mutants have presence of pistils in the tassel flowers (POPIT) and reduced tiller branch outgrowth. Genetic interactions between BR and GA biosynthetic mutants were developmentally specific and varied. Mutants in BR and GA were additive for plant height, indicating independent effects on this trait. BR mutants were epistatic to GA mutants for tiller outgrowth but GA mutants were epistatic to BR mutants for POPIT.

Specific and effective inhibition of GA and BR biosynthesis should recapitulate the phenotypes of these three differing genetic interactions seen in the mutants. Interpretation of inhibitor studies requires confidence in the specificity and amplitude of inhibition caused by chemical treatment. Inhibitors of plant hormone biosynthesis are widely used as plant growth regulators in industry and basic science to modify plant metabolism and phenotype. Triazole PGRs are widely used, cheap to produce, and effective on both plants and fungi. The chemical structures of three triazole inhibitors containing the 1,2,4-triazole ring used in this study are shown in Figure 1. PGR specificity within plants is known to be poor with wide impacts across P450s (Figure 2). It is currently unknown how wide these impacts actually are.^18^

**Figure 1.**
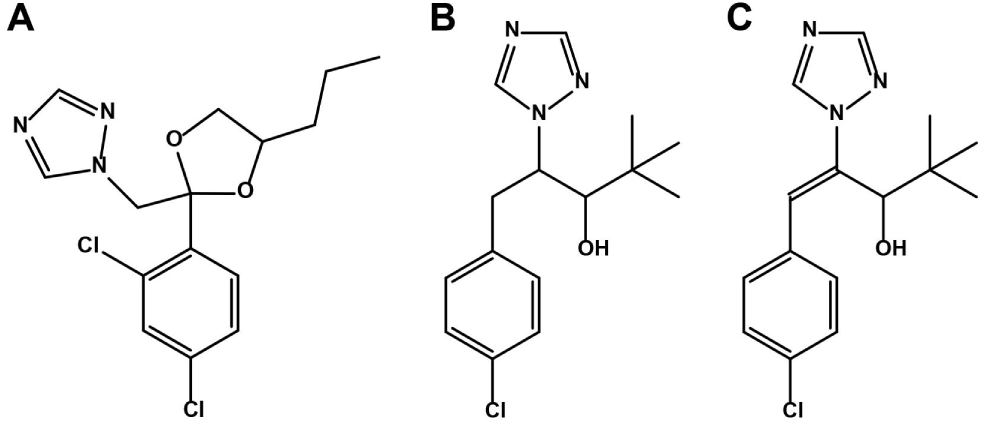
Structures of chemicals used in this study. Chemical structure of (A) propiconazole, (B) paclobutrazol, and (C) uniconazole.

**Figure 2.**
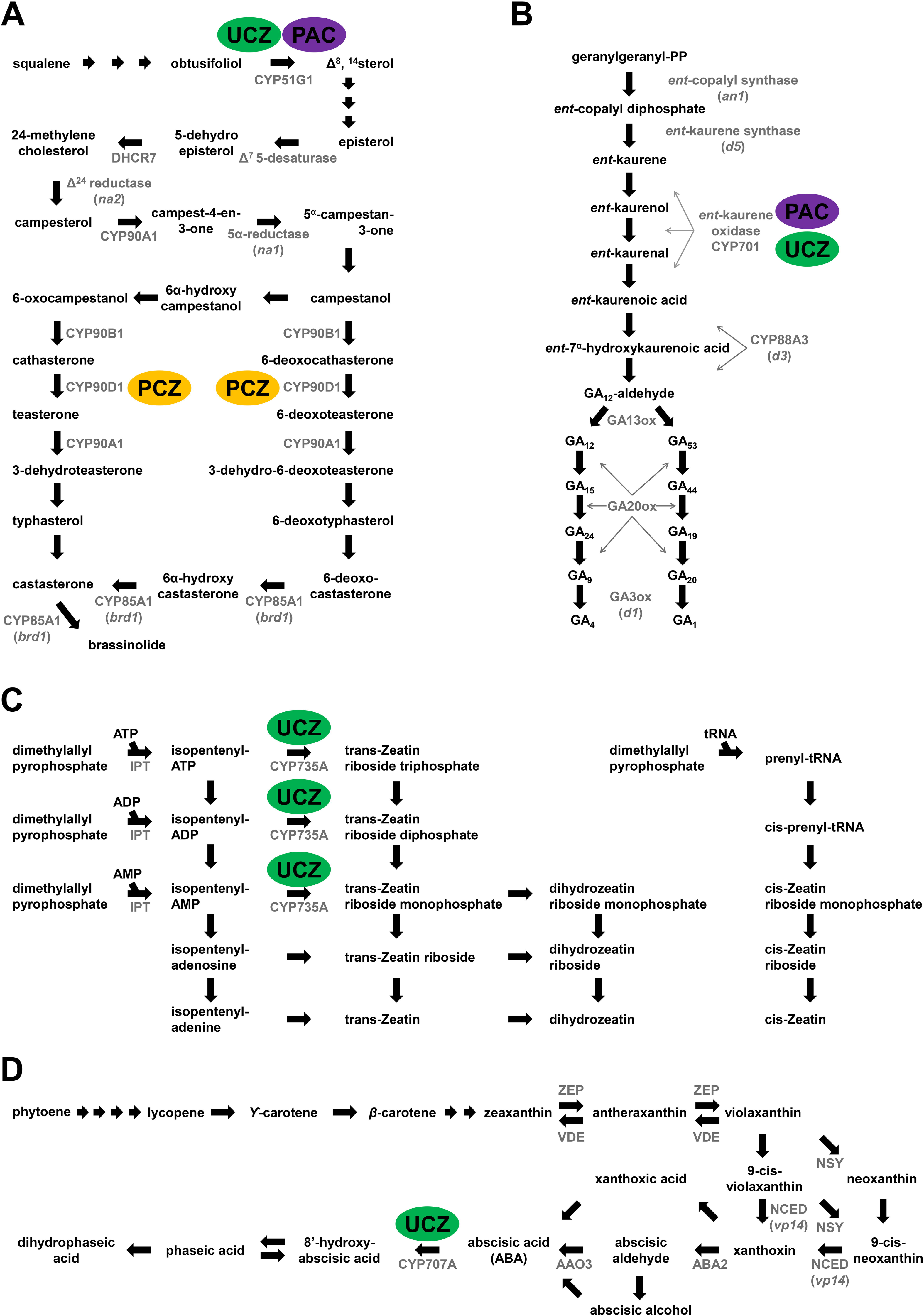
All three plant growth regulators are known to affect more than one pathway. (A) Brassinosteroid, (B) gibberellin, (C) cytokinin, and (D) ABA metabolic pathways. Metabolites are indicated in black text and enzymes by black arrows with respective names in grey text. Functionally characterized enzymes in maize are indicated in parentheses. Previously identified enzymes inhibited by PCZ (yellow), PAC (purple), and UCZ (green) are indicated by colored ovals.

Propiconazole (PCZ) was first identified as a fungal growth inhibitor that blocked the C-14 demethylation of lanosterol (Figure 1A).^19,20^ Propiconazole is a specific and potent inhibitor of BR biosynthesis, as evidenced by the reversal of growth inhibition by the co-application of BR.^21^ PCZ binds the CYP90D1 of Arabidopsis (Figure 2A) but it is unknown whether PCZ also binds other P450s in BR biosynthesis such as CYP90B1/DWF4, CYP90C1/ROT3, or CYP90A1/CPD.^22^ The related BR biosynthesis inhibitor, brassinazole, effectively binds to CYP90B1 and the effectiveness of these two inhibitors may stem from an ability to simultaneously inhibit multiple CYP90/CYP85 BR biosynthetic enzymes. Co-treatment of Arabidopsis with epi-brassinolide reversed the phenotypic effects of both brassinazole and PCZ, suggesting that the effects of these triazoles were predominantly due to BR inhibition.^21,23,24^ PCZ is also used as a fungicide in agriculture due to inhibition of ergosterol biosynthesis. It is possible that PCZ also affects structural sterol biosynthesis in plants. To date, no experiments have been conducted on PCZ and enzymes in other metabolic pathways to determine effects on other phytohormones or sterol biosynthesis.

Paclobutrazol (PAC) was also first identified as a fungicide (Figure 1B).^25^ In fungi, it inhibits C-14 demethylation, similar to PCZ. In barley and celery cell cultures it was shown to inhibit the 14α-demethylase CYP51 required for the conversion of obtusifoliol to Δ^8^-^14^sterol (Figure 2A).^26–28^ Supplementation of cell cultures with stigmasterol at high concentrations, or a mix of cholesterol and stigmasterol, reversed the growth retardation that resulted from PAC treatment of celery cell cultures. Ratios of stigmasterol to sitosterol, a 22-desaturation carried out by CYP710A, were also altered in PAC treated cells suggesting that this P450 may also be a target. The BR precursor campesterol was also reduced by PAC treatment, and it may be that PAC achieves growth retardation by affecting structural sterols as well as BR levels.^27^ PAC is also known to reduce GA levels by inhibiting the *ent*-kaurene oxidase/CYP701 in plants (Figure 2B).^18^ Hedden & Graebe^29^ showed the percent conversion of the CYP701 targets, *ent*-kaurene, *ent*-kaurenol, and *ent*-kaurenal, into later intermediates was greatly inhibited by 1 μM (2RS, 3RS)-PAC in a cell-free system from *Cucurbita maxima* endosperm. The different enantiomers of PAC have demonstrated differential effects on plants as compared to fungi. The 2R-3R diastereoisomer of PAC affected sterol biosynthesis in fungi but had no effect on plant height at 170.2 μM while the 2S 3S diastereoisomer dramatically reduced plant height at 34.0 μM.^25^ A number of other downstream effects have been documented in PAC-treated plants including reduced accumulation of 1-aminocyclopropane-1-carboxylic acid (ACC) and ethylene production in water-stressed apple seedlings, however the mode of action is unknown.^30^

The uniconazole (UCZ) structure differs from PAC by one desaturation leading to a double bond (Figure 1C) and has been shown to inhibit GA biosynthesis by affecting the same enzymes (Figure 2B).^18,31,32^ However, the height of a GA_3_-insensitive DELLA mutant in *Helianthus annuus* (sunflower) was further reduced by UCZ-treatment.^33^ UCZ has also been shown to affect structural sterol levels in *Oryza sativa* (rice) and pea (Figure 2A).^34,35^ In addition, UCZ treatment reduced castasterone accumulation in pea and blocked BR-induced tracheary element differentiation in *Zinnia elegans* L. mesophyll cells.^36,37^ Whether these phenotypes result from BR-inhibition, the dependence of BR-induced tracheary element differentiation on GA, or some other impact is unknown. Recently, it has been shown that UCZ affects ABA catabolism by inhibiting CYP707A in Arabidopsis and tobacco cell cultures (Figure 2D).^38–40^ UCZ also affects cytokinin biosynthesis; however, there are conflicting reports depending on the species tested. In Arabidopsis it inhibited trans-Zeatin accumulation (Figure 2C) but UCZ treatment in rice and *Glycine max* showed higher levels of trans-Zeatin.^41–43^ Izumi et al^41^ showed that ethylene levels were higher but that ABA was unaffected in UCZ-treated rice. Of the three PGRs used in this study, UCZ has been the most studied. The conflicting reports on hormonal levels in different species highlights the necessity of studying the specificity of PGRs within the species being tested due to potential differences in uptake, transport, or binding affinity by potential target P450s.^18^

We sought to test the model for BR and GA interaction in maize derived from genetics experiments.^17^ Given the multiple interactions between the mutants it is less likely that the same effects, and direction of interaction, will be generated by non-specific growth retardation or general toxicity of a chemical treatment. In addition, using the genetics experiments as orthogonal data confirming the utility of these inhibitors’ specificity in maize would permit stronger interpretation of inhibitor studies across the grasses, including species without the genetic resources of maize. Orthogonal data, if confirmatory, will also bolster our confidence in the interpretation of previous genetic interactions between BR and GA discovered in maize.^17^

We treated the *na2-1* BR- and *d5* GA-biosynthetic mutants with each of the three triazole inhibitors described above and measured phenotypes known to respond to BR and GA reduction. All three inhibitors were additive with the two mutants for plant height. UCZ had the strongest impact on plant height. Consistent with the expectation for reduced BR biosynthesis, PCZ enhanced the POPIT phenotype of BR mutants but was unable to induce POPIT in wild-type siblings or *d5* mutants at the concentrations employed. PCZ application phenocopied the BR-GA interaction and suppressed the outgrowth of tillers in *d5* mutants. Just like GA dwarfs, PAC treatment induced tillering in wild-type plants. Unlike GA dwarfs, PAC treatment did not rescue the POPIT phenotype of BR mutants nor did it induce anthers in ear florets. Only UCZ recapitulated all GA dwarf phenotypes including the effects on floral organ persistence and interactions with loss of BR biosynthesis, as UCZ treatment induced anther-ear in wild type and *na2-1*.

## Results

### Effect of hormone biosynthetic inhibitors on *na2-1* and wild-type siblings

As described previously,^17^ *na2-1* mutants overall height and organ lengths were shorter, lower leaves were more upright, mutants flowered later, and a majority of plants had POPIT. Mock-treated wild type and *na2-1* are shown in Figure 3 along with a PCZ-treated wild-type plant. PCZ treated wild types strongly resembled mock treated *na2-1* plants (Figure 3A-C). Treatment of *na2-1* plants with PCZ displayed an enhanced phenotype, indicating synergy between the loss of *na2* function and the effects of PCZ. Organs and internodes were shorter in PCZ-treated plants than in mock-treated plants for both *na2-1* and wild-type siblings (Table 1 and Figure 4A-B). PCZ treatment had a dramatic effect on *na2-1* mutant plant height indicating either bioactive BR accumulation in the mutant or non-specific inhibition of other growth-promoting metabolites by PCZ application. Primary tassel branch number was 10 fold lower in PCZ treated *na2-1* than in treated wild-type siblings, whereas mutants and wild types were indistinguishable in the mock treated samples (Table 1). Tassel length was also non-additively affected by PCZ application to *na2-1* mutants, causing a decrease by 40% in wild type and 80% in *na2-1* mutants. Just as *na2-1* mutants flowered later than wild types, PCZ treatment extended days to flowering in wild types. PCZ treatment of the mutants further enhanced this delay (Table 1). Yet, in neither comparisons between *na2-1* and wild type nor in PCZ treatments did the number of leaves per plant change. Taken together, PCZ reproduced all but the POPIT phenotype in wild-type treated plants and enhanced all loss-of-BR phenotypes in the *na2-1* mutant, including increased POPIT penetrance.

**Figure 3.**
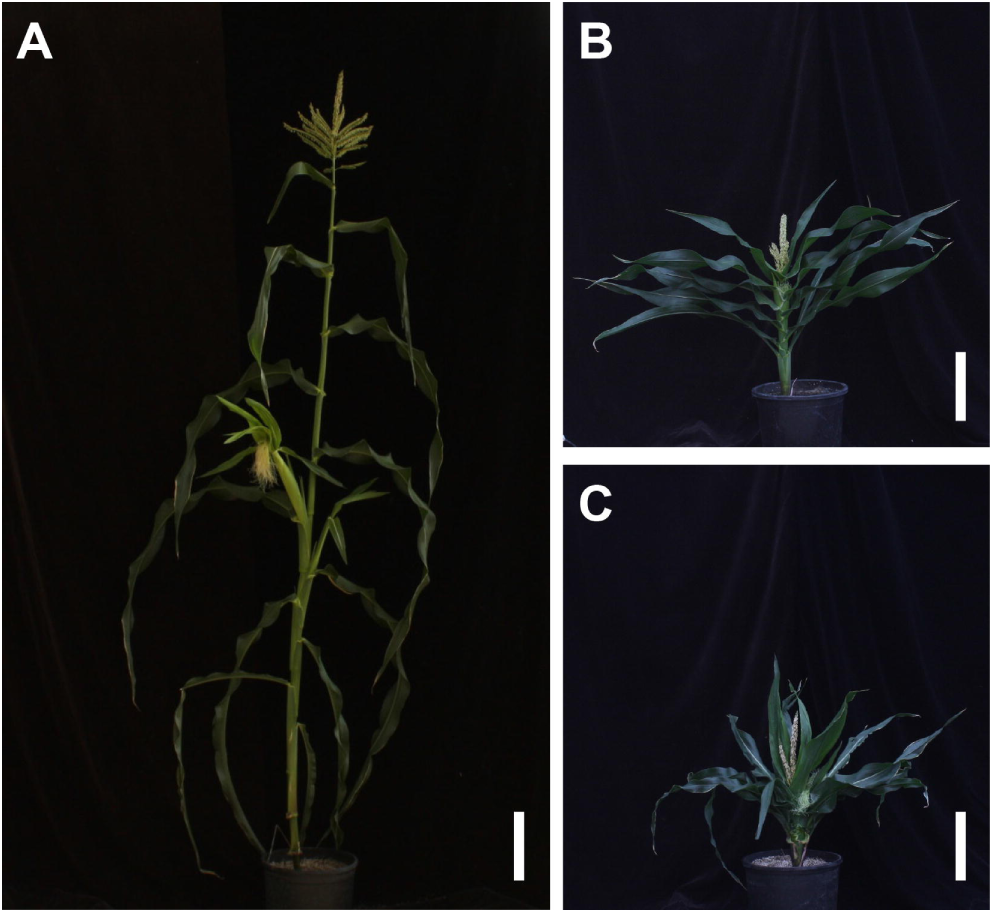
Effects of *na2-1* or PCZ on maize plant architecture. Mock treated (A) wild-type plant and (B) *na2-1* plant. (C) PCZ-treated wild-type plant.

**Figure 4.**
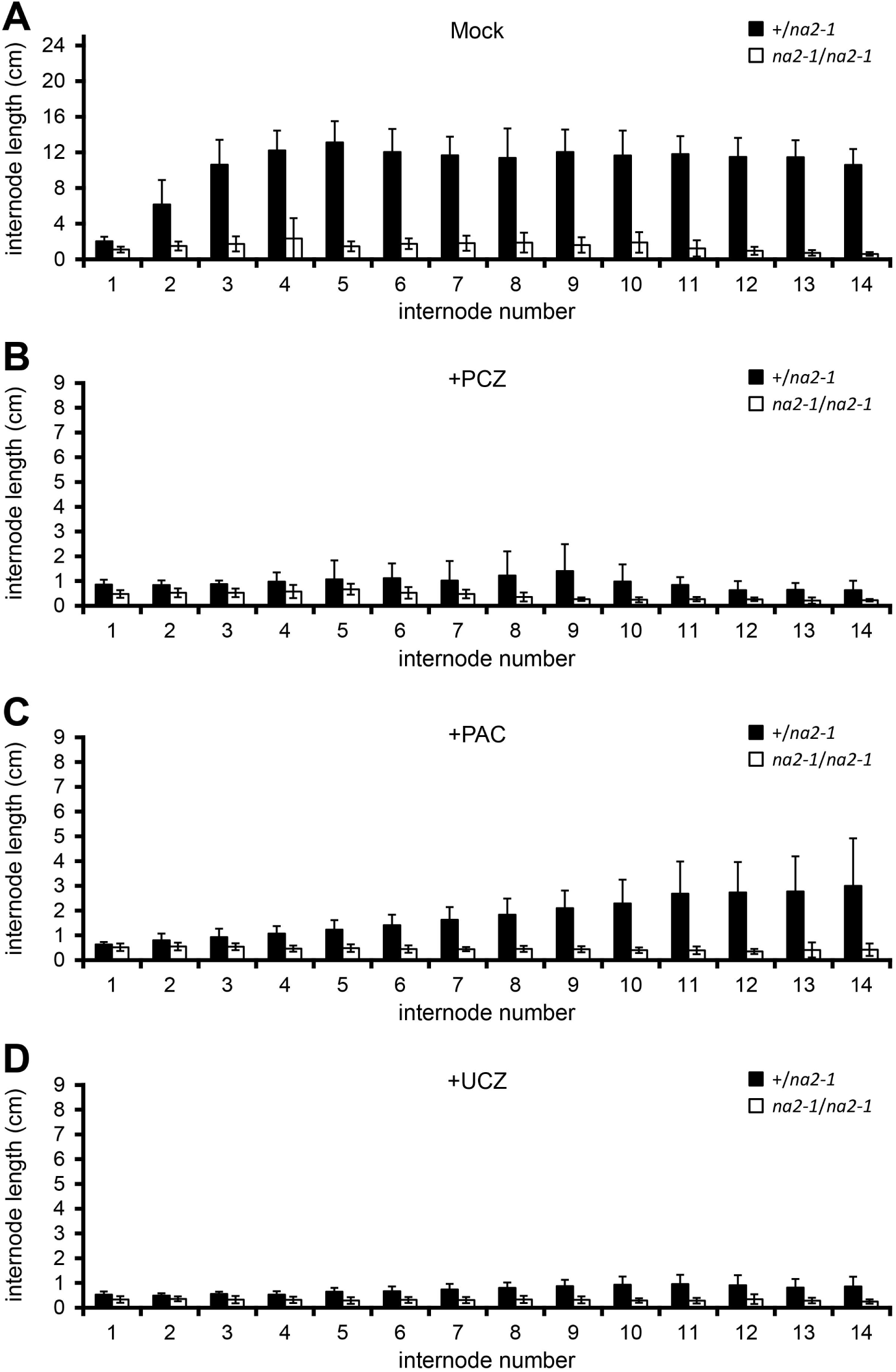
Effects of plant growth regulators and genotype on internode length. Mean internode lengths in cm and SD of (A) mock-treated, (B) PCZ-treated, (C) PAC-treated, and (D) UCZ-treated *na2-1* and wild-type siblings.

**Table 1.**
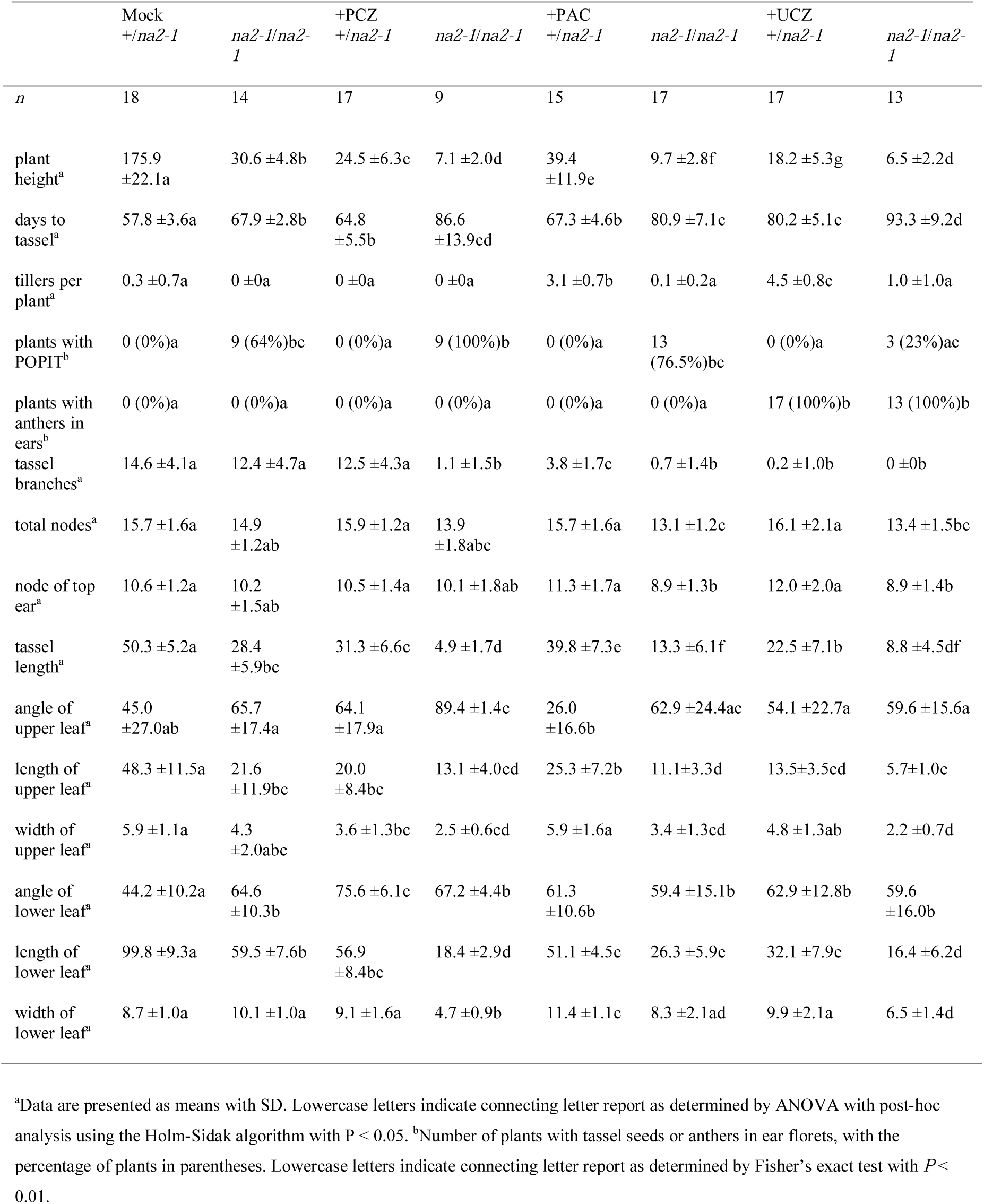
Morphometric analysis of *na2-1* and heterozygous wild-type siblings treated with PCZ, PAC, and UCZ.

|  | Mock<br>+/na2-1 | na2-1/na2-1 | +PCZ<br>+/na2-1 | na2-1/na2-1 | +PAC<br>+/na2-1 | na2-1/na2-1 | +UCZ<br>+/na2-1 | na2-1/na2-1 |
| --- | --- | --- | --- | --- | --- | --- | --- | --- |
| *n* | 18 | 14 | 17 | 9 | 15 | 17 | 17 | 13 |
| plant height<sup>a</sup> | 175.9<br>±22.1a | 30.6 ±4.8b | 24.5 ±6.3c | 7.1 ±2.0d | 39.4<br>±11.9e | 9.7 ±2.8f | 18.2 ±5.3g | 6.5 ±2.2d |
| days to tassel<sup>a</sup> | 57.8 ±3.6a | 67.9 ±2.8b | 64.8<br>±5.5b | 86.6<br>±13.9cd | 67.3 ±4.6b | 80.9 ±7.1c | 80.2 ±5.1c | 93.3 ±9.2d |
| tillers per plant<sup>a</sup> | 0.3 ±0.7a | 0 ±0a | 0 ±0a | 0 ±0a | 3.1 ±0.7b | 0.1 ±0.2a | 4.5 ±0.8c | 1.0 ±1.0a |
| plants with POPIT<sup>b</sup> | 0 (0%)a | 9 (64%)bc | 0 (0%)a | 9 (100%)b | 0 (0%)a | 13<br>(76.5%)bc | 0 (0%)a | 3 (23%)ac |
| plants with anthers in ears<sup>b</sup> | 0 (0%)a | 0 (0%)a | 0 (0%)a | 0 (0%)a | 0 (0%)a | 0 (0%)a | 17 (100%)b | 13 (100%)b |
| tassel branches<sup>a</sup> | 14.6 ±4.1a | 12.4 ±4.7a | 12.5 ±4.3a | 1.1 ±1.5b | 3.8 ±1.7c | 0.7 ±1.4b | 0.2 ±1.0b | 0 ±0b |
| total nodes<sup>a</sup> | 15.7 ±1.6a | 14.9<br>±1.2ab | 15.9 ±1.2a | 13.9<br>±1.8abc | 15.7 ±1.6a | 13.1 ±1.2c | 16.1 ±2.1a | 13.4 ±1.5bc |
| node of top ear<sup>a</sup> | 10.6 ±1.2a | 10.2<br>±1.5ab | 10.5 ±1.4a | 10.1 ±1.8ab | 11.3 ±1.7a | 8.9 ±1.3b | 12.0 ±2.0a | 8.9 ±1.4b |
| tassel length<sup>a</sup> | 50.3 ±5.2a | 28.4<br>±5.9bc | 31.3 ±6.6c | 4.9 ±1.7d | 39.8 ±7.3e | 13.3 ±6.1f | 22.5 ±7.1b | 8.8 ±4.5df |
| angle of upper leaf<sup>a</sup> | 45.0<br>±27.0ab | 65.7<br>±17.4a | 64.1<br>±17.9a | 89.4 ±1.4c | 26.0<br>±16.6b | 62.9 ±24.4ac | 54.1 ±22.7a | 59.6 ±15.6a |
| length of upper leaf<sup>a</sup> | 48.3 ±11.5a | 21.6<br>±11.9bc | 20.0<br>±8.4bc | 13.1 ±4.0cd | 25.3 ±7.2b | 11.1 ±3.3d | 13.5 ±3.5cd | 5.7 ±1.0e |
| width of upper leaf<sup>a</sup> | 5.9 ±1.1a | 4.3<br>±2.0abc | 3.6 ±1.3bc | 2.5 ±0.6cd | 5.9 ±1.6a | 3.4 ±1.3cd | 4.8 ±1.3ab | 2.2 ±0.7d |
| angle of lower leaf<sup>a</sup> | 44.2 ±10.2a | 64.6<br>±10.3b | 75.6 ±6.1c | 67.2 ±4.4b | 61.3<br>±10.6b | 59.4 ±15.1b | 62.9 ±12.8b | 59.6<br>±16.0b |
| length of lower leaf<sup>a</sup> | 99.8 ±9.3a | 59.5 ±7.6b | 56.9<br>±8.4bc | 18.4 ±2.9d | 51.1 ±4.5c | 26.3 ±5.9e | 32.1 ±7.9e | 16.4 ±6.2d |
| width of lower leaf<sup>a</sup> | 8.7 ±1.0a | 10.1 ±1.0a | 9.1 ±1.6a | 4.7 ±0.9b | 11.4 ±1.1c | 8.3 ±2.1ad | 9.9 ±2.1a | 6.5 ±1.4d |

<sup>a</sup>Data are presented as means with SD. Lowercase letters indicate connecting letter report as determined by ANOVA with post-hoc analysis using the Holm-Sidak algorithm with $P < 0.05$ . <sup>b</sup>Number of plants with tassel seeds or anthers in ear florets, with the percentage of plants in parentheses. Lowercase letters indicate connecting letter report as determined by Fisher's exact test with $P < 0.01$ .

As expected if PAC primarily acts via the inhibition of GA biosynthesis, all organs and overall plant height were reduced by PAC treatment in both *na2-1* and wild-type siblings (Table 1 and Figure 4A,C). Treatment of wild-type siblings with PAC resulted in a significant increase in tiller outgrowth similar to the GA biosynthetic mutant *d5*. The mock-treated *na2-1* mutants did not tiller and *na2-1* suppressed the tiller outgrowth affected by PAC-treatment. Thus, PAC treatment phenocopied the *na2-1/d5* double mutants, where *na2* was required for tillering induced by a loss of GA. Unlike *na2-1/d5* double mutants, PAC did not suppress POPIT in *na2-1* mutants indicating that PAC treatment was insufficient to cause all of the changes in plant growth affected by genetic loss of GA biosynthesis. Similar to the phenotypes of *d5* mutants,^17^ PAC treatment of wild-type siblings caused the lower leaf angle to become more upright. Contrastingly, the upper leaf angles of wild-type siblings went the opposite direction and were less upright following PAC treatment. As was the case for tillering, *na2-1* suppressed the effects of PAC on leaf angle and no discernable difference between mock and PAC treatments were observed. Similar to PCZ, primary tassel branch numbers of *na2-1* plants were reduced by PAC treatments. Flowering time was increased by PAC treatment of both *na2-1* and wild types. Surprisingly, PAC treatment reduced the number of leaves before the top ear and the total number of leaves per plant in *na2-1* but not wild-type siblings. This demonstrates that PAC reduced the plastochron index in *na2-1* mutants. For all phenotypes except suppression of POPIT and induction of anther-ear, PAC recapitulated the phenotypes observed in *d5* and *d1* mutants as well as the genetic interactions of *d5* and *na2*.

Treatment with UCZ reproduced all of the effects of a loss of GA biosynthesis and the genetic interactions between *d5* and *na2*. The effect of UCZ on organ length and plant height of *na2-1* and wild-type siblings was similar to what was observed in *na2-1/d5* double mutants, PAC, and PCZ treatments (Table 1 and Figure 4A, D). Consistent with a strong inhibition of GA biosynthesis, UCZ caused ears of both *na2-1* and wild-type siblings to exhibit florets with persistent anthers. Similar to *d5* mutants and PAC treatment, UCZ treated wild types profusely tillered. Similar to the *na2-1/d5* double mutants, and consistent with the genetic BR and GA interactions, *na2-1* strongly suppressed UCZ-induced tillering. UCZ treatment reduced the frequency of POPIT in *na2-1* by over two fold, similar to the suppression of POPIT in *na2-1/d5* double mutants, but this difference was not statistically significant from mock treated *na2-1* mutants. UCZ treatments had a synergistic effect on leaf width in combination with *na2-1* and caused *na2-1* leaves to become narrower. UCZ resulted in a dramatic decrease in primary tassel branch number, similar to the other inhibitors. UCZ treatment delayed the number of days before tassel emergence in both mutants and wild-type siblings. Similar to PAC treatment, both the number of leaves before the top ear and the total number of leaves per plant were dramatically reduced in *na2-1*, but not wild-type siblings, by UCZ treatment. Thus, for UCZ treatments this extreme delay without additional production of leaves resulted in an increase in plastochron index in *na2-1* UCZ treated plants. Taken together, UCZ treatments reproduced all of the phenotypes present in GA biosynthetic mutants and consistently affected the same interactions with loss of BR biosynthesis observed in *na2-1/d5* double mutants.

### Effect of hormone biosynthetic inhibitors on ***d5*** and wild-type siblings

Similar to what was found previously,^17^ *d5* mutants flowered later, were shorter, had shorter organs, exhibited persistent anthers in the ear florets, had reduced primary tassel branch numbers, and tillered profusely. Shown in Figure 5 are mock-treated wild-type and *d5* plants compared to PAC-treated and UCZ-treated wild-type siblings. Similar to what was observed for the wild-type siblings of *na2-1*, PAC treatment largely recapitulated the phenotypes affected by loss of function mutants in GA biosynthesis (Figure 5A-C). At the concentrations employed, one out of sixteen wild types exhibited anther-ear and the reduction in tassel length was less in PAC treated wild types than in mock-treated *d5* mutants. PAC further reduced the height and organ lengths of *d5* plants (Table 2 and Figure 6A, C). Similar to the effects of PAC on primary tassel branch number in *na2-1* mutants, PAC reduced primary tassel branch number to zero in *d5*. The *d5* mutants had greater leaf widths than wild-type controls. PAC treatment increased leaf widths of wild types but PAC treated *d5* mutants had narrower leaves than mock treated mutants. The *d5* mutants had the same number of nodes and, unlike PAC-treated *na2-1*, treatment of *d5* mutants with PAC did not affect node number. PAC treatment delayed flowering in wild-type siblings and further delayed flowering in *d5* mutants as compared to mock treatments. Thus, a delay without increased leaf numbers suggests an increase in plastochron index, though not as dramatic as the synergistic effects of inhibitor treatment in *na2-1* mutants. All PAC treated phenotypes were consistent with a loss of GA, but the low frequency of anther-ear, similar to the lack of suppression of POPIT in *na2-1*, suggest that at this concentration PAC is a weak GA inhibitor in maize.

**Figure 5.**
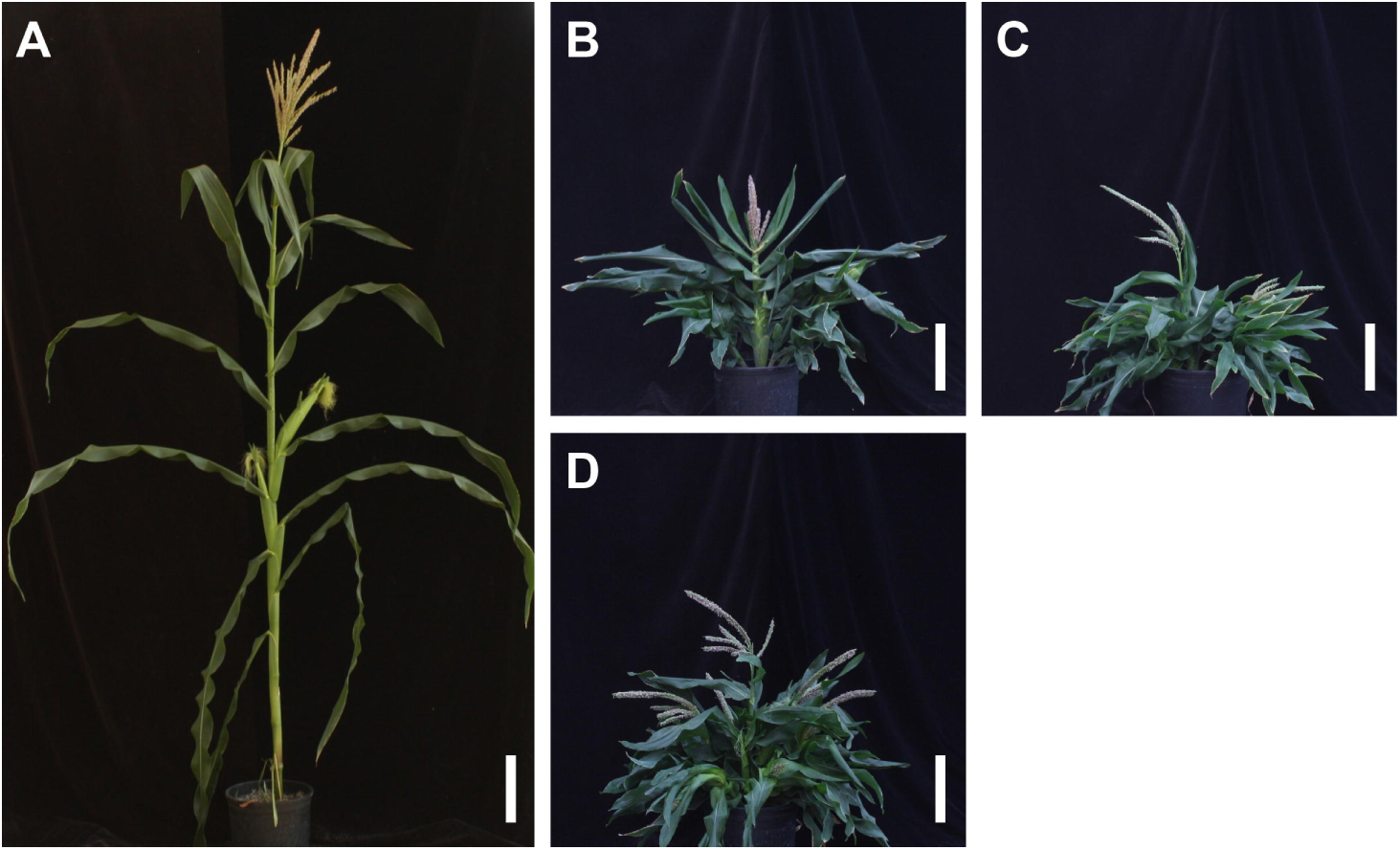
Effects of *d5*, PAC, and UCZ on maize plant architecture. Mock treated (A) wild-type plant and (B) *d5* plant. (C) PAC-treated and (D) UCZ-treated wild-type plant.

**Table 2.** Morphometric analysis of *d5* treated with PCZ, PAC, and UCZ.

|  | Mock |  | +PCZ |  | +PAC |  | +UCZ |  |
| --- | --- | --- | --- | --- | --- | --- | --- | --- |
|  | *+/d5* | *d5/d5* | *+/d5* | *d5/d5* | *+/d5* | *d5/d5* | *+/d5* | *d5/d5* |
| *n* | 16 | 16 | 16 | 16 | 16 | 16 | 16 | 16 |
| plant height<sup>a</sup> | 196.3±17.8a | 33.8±7.3b | 26.4±5.2c | 12.2±2.7d | 45.5±12.8e | 10.1±1.9f | 26.4±7.1c | 6.5±1.3g |
| days to tassel<sup>a</sup> | 52.5±4.4a | 63.3±4.0bc | 58.3±5.8d | 72.9±5.6e | 60.0±5.5bd | 78.8±6.0e | 66.4±4.5c | 78.3±13.5e |
| tillers per plant<sup>a</sup> | 0.3±0.4ab | 3.1±1.4cd | 0±0a | 0.6±0.6b | 2.9±1.2c | 3.1±1.5c | 5.2±2.3d | 3.9±1.7cd |
| plants with POPIT<sup>b</sup> | 0 (0%)a | 0 (0%)a | 0 (0%)a | 0 (0%)a | 0 (0%)a | 0 (0%)a | 0 (0%)a | 0 (0%)a |
| plants with anthers in ears<sup>b</sup> | 0 (0%)a | 16 (100%)b | 0 (0%)a | 15 (94%)b | 1 (6%)a | 16 (100%)b | 16 (100%)b | 16 (100%)b |
| tassel branches<sup>a</sup> | 23.9±6.1a | 8.9±4.1b | 15.8±7.8c | 3.8±3.0d | 5.1±4.7d | 0±0e | 1.6±1.5f | 0±0e |
| total nodes<sup>a</sup> | 14.1±1.2a | 14.4±2.1a | 14.2±0.8a | 14.9±2.0a | 14.7±1.2a | 15.0±2.3a | 14.9±1.7a | 13.9±2.2a |
| node of top ear<sup>a</sup> | 8.6±1.3a | 8.5±1.3a | 9.6±1.1a | 9.3±1.8a | 9.8±0.8a | 8.4±2.3a | 9.5±2.0a | 7.8±2.5a |
| tassel length<sup>a</sup> | 63.3±4.7a | 30.5±2.1b | 41.2±12.9c | 21.4±3.7d | 55.5±7.0e | 15.1±4.4f | 37.3±6.9c | 10.8±3.4g |
| angle of upper leaf<sup>a</sup> | 52.0±23.2abc | 39.7±16.3a | 49.7±17.2ab | 74.4±7.7d | 46.3±14.2a | 66.9±16.9bcd | 51.6±25.0abc | 72.5±12.9cd |
| length of upper leaf<sup>a</sup> | 53.2±7.4a | 28.9±6.5bc | 29.4±10.8bc | 19.6±5.2d | 34.5±6.2b | 10.0±2.5e | 22.6±6.2cd | 7.0±2.3f |
| width of upper leaf<sup>a</sup> | 7.2±1.0abc | 7.7±1.6ab | 5.9±1.5c | 6.5±1.3bc | 8.0±1.0a | 4.0±1.3d | 6.2±1.3bc | 3.6±1.0d |
| angle of<br>lower leaf<sup>a</sup> | 50.0±10.0a | 63.8±6.5b | 70.6±8.1b | 71.6±8.3b | 68.1±6.8b | 69.7±7.6b | 68.1±7.0b | 68.8±11.2b |
| length of<br>lower leaf<sup>a</sup> | 107.9±8.7a | 56.5±8.0b | 57.3±9.4b | 34.7±7.2c | 52.3±9.3b | 16.3±2.0d | 34.4±7.1c | 11.9±1.7e |
| width of<br>lower leaf<sup>a</sup> | 7.8±0.8a | 10.0±1.3bc | 9.8±0.8bc | 10.2±1.4bc | 10.6±0.9c | 7.3±1.0a | 9.2±1.4b | 5.9±1.1d |

<sup>a</sup>Data are presented as means with SD. Lowercase letters indicate connecting letter report as determined by ANOVA with post-hoc analysis using the Holm-Sidak algorithm with $P < 0.05$ . <sup>b</sup>Number of plants with tassel seeds or anthers in ear florets, with the percentage of plants in parentheses. Lowercase letters indicate connecting letter report as determined by Fisher's exact test with $P < 0.01$ .

**Figure 6.**
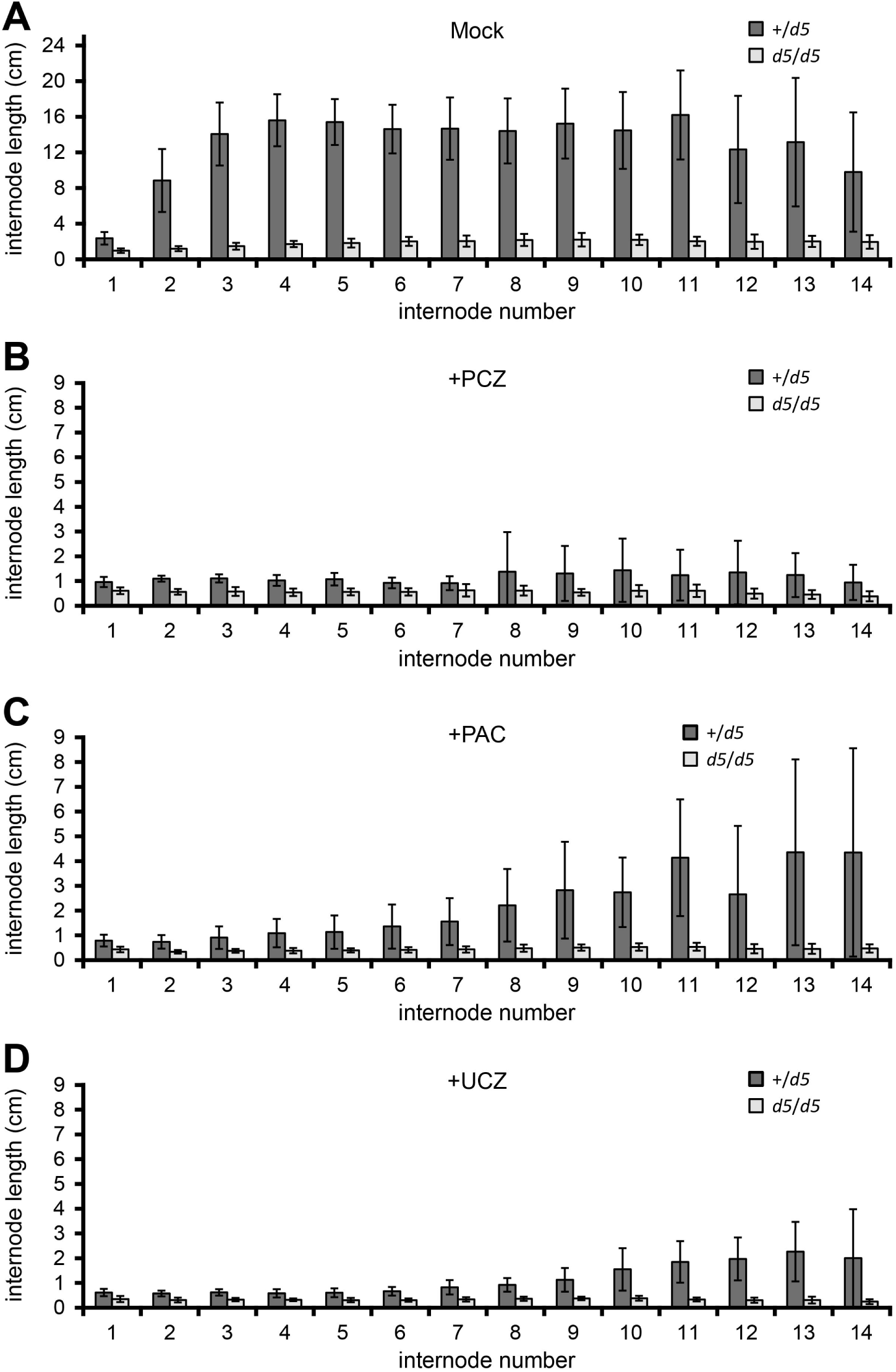
Effects of plant growth regulators and genotype on internode length. Mean internode lengths in cm and SD of (A) mock-treated, (B) PCZ-treated, (C) PAC-treated, and (D) UCZ-treated *d5* and wild-type siblings.

As was observed in applications of UCZ to *na2-1* and wild types, application of UCZ to *d5* and wild-type siblings recapitulated all loss-of-GA phenotypes (Figure 5A-B, D). Application of UCZ decreased plant height and the lengths of all organs and internodes for both *d5* and wild-type siblings (Table 2 and Figure 6A, D). Similar to the effects of UCZ on *na2-1*, UCZ narrowed both the upper and lower leaves of *d5* mutants. Unlike PAC treatment, UCZ induced anthers in the ear florets of all wild-type siblings, consistent with it effectively inhibiting GA biosynthesis at the concentration employed. UCZ substantially reduced primary tassel branch number in *d5* mutants, just as it had for *na2-1*. While not affecting an increase in the number of leaves produced per plants, UCZ treatment increased days to tassel emergence for both *d5* and wild-type siblings. Thus, UCZ also appears to increase flowering time though no synergism affecting node number was observed between *d5* and UCZ, consistent with the mode of action affecting the same pathway.

The BR inhibitor PCZ reduced overall plant height, internode lengths, and the lengths of all measured organs for both *d5* and wild-type siblings (Table 2 and Figure 6A-B). Primary tassel branching decreased dramatically in both PCZ treated *d5* mutants and their wild-type siblings. PCZ application at this concentration did not reproduce the *na2-1* effect on POPIT in the wild-type controls and as expected from the *d5* suppression of POPIT in *na2-1/d5* double mutants no POPIT was observed in PCZ treated *d5* plants.^17^ Consistent with the inability of *na2-1* to suppress anther-ear in *d5*,^17^ PCZ treatment failed to suppress persistence of anthers in ear florets of *d5* mutants. The *d5* mutants exhibited increased tillering, which we previously demonstrated was dependent on BR.^17^ PCZ treatment suppressed tiller outgrowth induced by the loss of *d5*, just as *na2-1* suppressed tiller outgrowth in *na2-1/d5* double mutants.^17^ PCZ treatment increased the angle of the upper leaf in *d5* mutants and the angle of the lower leaf in wild types. Days to tassel emergence were greater in PCZ treated *d5* and wild-type siblings consistent with the flowering delay in double mutants and PAC treatment of *na2-1* plants. Thus, PCZ treatment of *d5* mutants and wild-type siblings delayed flowering, but no discernable difference was observed in the total number of nodes. Different from treatment of *na2-1* treated with PAC, synergism was not detected when *d5* was combined with PCZ treatment at this concentration for effect of total node number.

## Discussion

We report here the effects of PCZ, PAC, and UCZ treatment on the phenotypes of *na2-1, d5*, and wild-type siblings at maturity. PCZ-treatment of wild-type plants was able to recapitulate the phenotype of *na2-1*, except for the POPIT phenotype (Table 1 and Figure 3B-C). However, PCZ at twice the concentration (500 μM) used in our study induced POPIT in wild-type siblings of the *na1-1* mutant, which was in a unknown mixed-parentage Mu-active genetic background.^8^ Off-target and non-specific effects of inhibitors are a greater concern as concentrations go up and we sought to minimize these with lower concentrations. PCZ-treatment did decrease plant height and increase POPIT in *na2-1* (Table 1). These results, and our unpublished observations that *brd1* mutants and *na1/na2* double mutants are more severely dwarfed than *na1* or *na2* single mutants suggests that disruption of this gene may not be a complete knock-out of brassinosteroid biosynthesis. Alternative bioactive BRs can accumulate in rice brassinosteroid biosynthetic mutants due to the existence of a bypass pathway to C29/28 and C27 bioactive BRs and the same alternative BR pathway may exist in maize.^44^ Targeted metabolic analysis of these alternative BRs in the maize mutants is required to determine if this is the case.

UCZ treatment to wild-type plants was able to phenocopy *d5* mock-treated plants and UCZ treated *na2-1* plants displayed all the interactions predicted for loss of both GA and BR biosynthesis from *na2-1/d5* double mutant studies (Table 1 and 2). UCZ reduced plant height, promoted tiller outgrowth, induced anther persistence in the ear, and suppressed the *na2-1* mutant dependent POPIT (Table 1 and 2). These phenotypic results are consistent with UCZ affecting maize growth via specific inhibition of GA biosynthesis at the concentrations employed. However, the suppression of tiller formation induced by a loss of *na2* was the least effective of all treatment and genetic combinations observed (Table 1). Our experiments do not rule out inhibition of P450 enzymes outside of GA biosynthesis. Indeed it was established that UCZ inhibits ABA catabolism in Arabidopsis at concentrations lower than employed here and that specific inhibition of recombinant CYP707A occurs in with an IC50 of 68nM (Figure 2D).^45^ This reaction contributes to the formation of phaseic acid which has been demonstrated to have additional biological activities of its own (Figure 2D).^46,47^ The perfect agreement of the developmental phenotypes caused by UCZ and the GA mutants was somewhat surprising in light of these additional modes of action (Figure 2A-D). It is possible that the weak suppression of tiller formation by *na2-1* in UCZ treatments may result from inhibition of other P450s. It is formally possible, though we have no evidence indicating this is the case, that UCZ might inhibit P450s in strigolactone biosynthesis or that ABA might enhance tillering in maize. The simplest interpretation of our results is that UCZ affects these changes in development via inhibition of GA, for which there is ample evidence. ^18^ Further research, particularly work comprehensively evaluating the binding and catalytic inhibition by these compounds on isolated or recombinant P450s, as was done for ABA catabolism and UCZ, is needed to test these hypotheses and establish the specificity of all triazole inhibitors.

PAC treatment was unable to phenocopy all of the effects of GA deficient mutants in maize. PAC treatment of wild type and *na2-1* did not promote anther retention in the ear florets nor did it suppress POPIT in *na2-1* plants (Table 1). This contrasts with both UCZ treatments and our previous genetic results.^17^ The concentration of PAC used was the same as UCZ, but it may be that PAC is a weaker inhibitor as has been demonstrated in sunflower, *Impatiens wallerana, Salvia splendens, Tagetes erecta* L., and *Petunia hybrid*.^48,49^ If PAC is effectively inhibiting GA biosynthesis, this would contradict maize double mutant analyses where GA biosynthetic mutants were epistatic to BR biosynthetic mutants for the expression of POPIT.^17^ More likely the discordance between PAC and genetic ablation resulted from incomplete inhibition of GA biosynthesis by our repeated root drenches with 60 μM PAC. The effects of treatment on height and stimulation of tiller outgrowth in wild-type plants, and suppression of outgrowth by *na2-1*, matched the effects of reduced GA biosynthesis (Table 1 and 2). The inability of PAC treatment to induce anther-ear in any genotype, except for *d5*, or suppress POPIT in *na2-1* may be due to the concentration used, unspecific inhibition of other pathways besides GA biosynthesis (Figure 2), or combination of the two. The observation that PAC growth arrest can be reversed by application of sterols raises the possibility that PAC achieves some height reduction by affecting both BR and GA levels.^27^ If BR biosynthesis is inhibited by PAC treatment this could increase the penetrance of POPIT in the *na2-1* mutant, as was observed in PCZ treatment, and require greater reduction in GA to prevent floral organ persistence. However, the lack of anthers in ear florets in PAC treatments, which were observed in all maize GA mutants, suggested that PAC might simply be a weaker GA biosynthesis inhibitor.

One unexpected finding was the effect of GA biosynthesis on tassel branch numbers. The *d5* mutants had fewer primary tassel branches than wild-type siblings but *na2-1* mutant primary tassel branch numbers were unaffected in both the results of this study and our previous work (Table1 & Table 2).^17^ Treatment of wild-type plants with PAC or UCZ also reduced primary tassel branch numbers. Unlike the *na2-1* mutants, PCZ treatment reduced tassel branch number, however the interpretation of these results is complicated by the potential for genetic background effects. The wild-type background of the *d5* mutants, but not the wild-type siblings of *na2-1*, had fewer primary tassel branches following PCZ treatment. The *na2-1* mutants did not have fewer branches in mock treatments but PCZ treatment of mutants did reduce tassel branch numbers. The inability of PCZ to reduce tassel branching in *na2-1* wild-type siblings casts some doubt into whether the full tassel branching phenotype of *na2-1* might be different in the genetic background of our *d5* mutants. Residual BR biosynthetic activity in all *na2* alleles is suggested by the stronger phenotype of *brd1* mutants and *na1/na2* double mutants (Dilkes and Best, data not shown) and it may be that the residual BRs are sufficient to maintain normal tassel branching in *na2-1* mutants.^16^ Alternatively, PCZ might have effects on regulators of tassel branching encoded by P450s outside of BR biosynthesis. All triazoles affected branching in *na2-1* and *d5* mutants. In our previous work, increasing GA by exogenous application to *na2-1, d5*, and wild type growing apices did not alter tassel branch numbers.^17^ Additional work determining the physiological basis of triazole application on tassel branching may uncover as yet unknown impacts of GA in tassel architecture, or implicate triazoles in the inhibition of other branch-promoting signals.

We repeatedly treated plants with PGRs and this continuous treatment was required to observe the dramatic decrease of plant height. This requirement suggests that maize can catabolize or inactivate these PGRs over the course of a few days. Reduction in the concentrations of bioactive PGRs should permit the biosynthesis of phytohormones to return and promote growth in responsive cells. We ceased treatment at tassel emergence during the elongation of the uppermost internodes and the lengths of these internodes were not as affected in our treated plants (Figure 4). We propose that PGR inactivation rather than phytohormone-independent growth underlies these differences in internode responses.

Days to tassel emergence was significantly longer for *na2-1* and *d5* (Tables 1 and 2) compared to their wild-type siblings similar to our earlier work.^17^ Wild-type siblings treated with PCZ, PAC, or UCZ had greater days to tassel emergence than controls. Every mutant-triazole combination further increased days to tassel emergence. When the *na2-1* mutant was combined with PAC or UCZ treatment, this resulted in a synthetic interaction producing fewer nodes even though plants took 2-3 weeks longer to flower. The plastochron index is the rate of lateral organ initiation from the meristem. We quantified total number of nodes and time to tassel emergence, so it is possible that we could overestimate the effect of these treatments as decreased elongation may delay emergence by a few days. However, the average increase of days to tassel emergence of *na2-1* or *d5* treated with any one inhibitor was an average of 19.0 days or 13.4 days, respectively. These results could be due to synergistic effects between BR and GA or inhibition of another cytochrome P450 by the triazoles. Overexpression of the maize *plastochron1* gene, which encodes cytochrome P450 CYP78A1, increases leaf number but decreased plastochron index.^50^ Loss of function of CYP78A1 exhibited slower leaf elongation, but interpretation is limited by the presence of multiple CYP78A paralogs in the maize genome.

The ability to combine chemical and genetic inhibition of phytohormone biosynthesis to assess interactions and combinatorial effects is limited only by resource availability. The discovery of compounds with additional modes of action would enable work on the effects of all phytohormones on plant development. An additional triazole was identified by Ito et al^51^ that inhibits strigolactone (SL) biosynthesis. This inhibitor, TIS13, effectively reduced SL levels in roots and root exudates of rice and increased tiller outgrowth of the second tiller. In these tests, PAC and UCZ had no effect on SL levels in root or root exudates, eliminating off-target inhibition of SL biosynthesis as a possible side-effect of PAC and UCZ treatments in rice. SLs control tiller outgrowth in maize.^52^ If SL also affects tassel branching, we would expect treatment of maize with TIS13 or GR24, a synthetic SL, to cause a decrease or increase of primary tassel branching, respectively. It may be that loss of GA affects tillering and reduced tassel branching via SL accumulation or signaling. A similar study to this one, using TIS3 application to GA and BR biosynthetic mutants of maize as well as UCZ, PCZ, and PAC treatment of the *carotenoid cleavage dioxygenase8* mutant of maize should provide a test of this hypothesis.^52^ Similarly, identification of specific inhibitors of cytochrome P450s in other metabolic pathways would permit combinatorial assessment with available mutants to explore metabolite function in maize.

In conclusion, PCZ treatment of wild-type siblings largely phenocopied *na2-1* mutants and PAC or UCZ treatment predominantly phenocopied *d5* mutants. PAC did not affect sexual organ persistence, unlike GA biosynthetic mutants or UCZ treatment. There was no strong evidence of non-specific growth inhibition by these three triazole inhibitors in maize. Unexpectedly, primary tassel branch number was reduced in *na2-1* or *d5* treated with PCZ, PAC, or UCZ. Thus, either all three triazoles another pathway controlling tassel branching or BR and GA biosynthetic pathways interact to affect tassel branching. Plastochron index was synergistically increased by PCZ, PAC, or UCZ treatment of both *na2-1* and *d5*. The mechanism for this is unknown, and we did not measure flowering time in *na2-1/d5* double mutants in our previous study.^17^ The similarity of the effects on plant height, tiller outgrowth, and floral organ persistence in the mutant analyses and inhibitor treatments were consistent with the genetic model proposed previously.^17^ The concordance of phenotypes increased the confidence of interpretation for UCZ and PCZ as GA and BR inhibitors, respectively. Using these inhibitors in the greenhouse via simple repeated pot drench will allow for rapid assessment of phytohormone function, even in the absence of genetic resources. Confidence in the interpretation of phenotypes resulting from inhibitor application should encourage exploration of BR and GA effects on growth and development in grass species such as *Sorghum bicolor, Panicum virgatum, Saccharum officinarum, Miscanthus sinensis*, and *Setaria virdis*.

## Materials and Methods

### Plant growth and genetic materials

Maize was planted in the winter of 2013 at the Purdue Horticulture Plant Growth Facility in 2 gallon pots with 2:1 mixture of Turface MVP:peat germinating mix and irrigated with fertilizer (MiracleGro Exel at 200ppm). Supplemental light was provided by sodium lamps for a L:D cycle of 16:8 with target temperatures of 27 °C Day 21°C Nighttime. All experiments were planted as complete randomized blocks, controlling for genotype and treatment. Maize genotypes *na2-1, d5*, and their respective wild types were described previously.^17^ Both mutants are recovered from segregating families. As such all comparisons are between mutants and wild-type siblings.

### Inhibitor treatments

All inhibitors were supplied to plants as a soil drench. Inhibitors were dissolved in methanol and added to irrigant to a final concentration of 0.15% methanol, 250 μM propiconazole (94% purity; ORICO Global, Zhuhai, China), 60 μM uniconazole (95% purity; Seven Continent, Zhangjiagang, China), or 60 μM paclobutrazol (0.4%; formulated as *Bonzi*, Syngenta, Basel, Switzerland). Inhibitor concentrations were chosen based on soilless media inhibition of efficacy and to closely phenocopy respective genetic mutants.^53^ A mock treatment consisting of 0.15% methanol was used as the control for all experiments. Plants were treated first at 10 days after germination (DAG) and on every subsequent 3^rd^ or 4^th^ day until tassel emergence. At the time of tassel emergence, treatments were stopped.

### Phenotypic measurements and statistical analysis

In total, 256 plants were phenotyped for an array of growth and developmental parameters. Plant height was measured as the distance from the soil to the uppermost leaf collar. The width at the widest point and the length from the leaf collar to the tip of the blade was determined for the second uppermost leaf (one below the flag leaf) and the leaf below the uppermost ear. These are described as the upper leaf and lower leaf in the tables. Leaf angle was measured as the difference from a 90° projection from the stem and the abaxial side of the leaf; greater values indicate a more upright leaf. The number of nodes on each plant, the length of each node, the node of the uppermost ear, the number of tassel branches, and number of tillers per plant were all recorded at maturity after stripping the plant of leaves. Tassels were inspected for POPIT and given a binary yes/no score. Ears were inspected for anthers and given a binary yes/no score. Flowering time was measured as DAG to visible tassel emergence from the leaf whorl. Statistical analysis of differences between genotypes and between treatments for continuously variable traits were performed by ANOVA with post-hoc pairwise tests by the Holm-Sidak method implemented in Daniel’s XL Toolbox add-in (ver. 7.2.6, http://xltoolbox.sourceforge.net) for Microsoft Excel (Redmond, WA). For the binary traits, differences were tested by Fisher’s exact tests using a Bonferroni correction for multiple tests that sets a corrected p<0.05 at the nominal p<0.002 as implemented in the Excel add-in Real Statistics Resource Pack (ver. 3.3.1, http://real-statistics.com).^54^

## Acknowledgments

We thank Rob Eddy and the Purdue Horticultural Plant Growth Facility team for the creation of superior growth environments. We also thank Jim Beaty and the Purdue University Agronomy Center for Research and Education (ACRE) for expert assistance in production of plant material. Johal laboratory members A. DeLeon, R. Zhan, and K. Chu helped with chemical treatments. N.B.B. would also like to thank M. Tuinstra and M. Hasegawa for advice and consultation. The Purdue Center for Plant Biology is gratefully acknowledged for their support and stimulation of intellectual rigor. This paper is offered in memory of Sewall Wright for his stunning breakthrough in biochemical genetics one century ago.

